# The Semantic Underpinnings of Speech Disorganization in Schizophrenia

**DOI:** 10.1101/2025.08.10.669589

**Authors:** Isaac Fradkin, Rani Moran, Rick A. Adams, Elizabeth Jefferies, Raymond J. Dolan

## Abstract

Effective communication relies on shared semantic representations and their context-sensitive retrieval. Accordingly, difficulties in communicating coherently, as seen in some patients with schizophrenia, may reflect impairments at the level of semantic structure or the retrieval process operating on this structure. Disentangling these components is challenging. Here, we address this problem using a word association task and magnetoencephalography (MEG), allowing us to examine behavioral markers of semantic structure and retrieval as well as semantic representation in the brain. For the latter, we detail an approach that contrasts the brain’s encoding of concepts as predicted by generic versus personalized semantic models. In so doing, we provide converging evidence for semantic structure idiosyncrasy in patients with schizophrenia who also exhibit speech disorganization. The findings advance our understanding of the semantic underpinnings of speech disorganization in schizophrenia, while also revealing the potential of using personalized semantic models to explain neural representation and behavior.

---

Thoughts are linked to one another in an associative manner. Virtually all models of semantic memory emphasize this associative nature^1–3^, implying that ideas and concepts trigger one another as a function of how closely they are connected. Yet, characterizing memory *structure* based on *retrieved output* is not trivial, not least because associations that come to mind are influenced by *how* people search within a given semantic structure^1,4–10^. Although semantic structure is assumed to be relatively stable, its efficient use necessitates a flexible adaptation to context, goals, and task-demands (i.e., *semantic control*)^2,11,12^. The importance of dissociating structure and process in semantic memory is widely acknowledged^1,2,4,7,11,12^, with implications for understanding creativity^13^, language, as well as neuropsychiatric syndromes thought to involve alterations in semantic memory^10,14–18^. Indeed, this distinction between structure and process is most relevant for clinical contexts, where the stability of (an atypical) semantic structure may present an obstacle to effective interventions^19–21^.

Here we focus on impairments in semantic cognition in schizophrenia, specifically with respect to their contribution to speech disorganization. In schizophrenia, previous studies have highlighted abnormalities either in semantic structure^10,16^ or at the level of controlled retrieval and semantic search processes^17,22^. Furthermore, whereas up to 50% of patients with schizophrenia exhibit incoherence and disorganization in speech, a phenomenon known as *Formal Thought Disorder* (Box 1)^23^, how putative underlying semantic impairments contribute to speech disorganization remains unknown.

## Box 1. Speech disorganization in formal thought disorder

Formal thought disorder involves different linguistic markers that render speech incoherent, idiosyncratic, and difficult to understand^23^. At the level of speech output, this involves ideas relating obliquely to one another (Derailment) or drifting away from the original question or response (Tangentiality). Inappropriate or made-up words can also be present (Paraphrasia and Neologisms, respectively). These characteristics, collectively known as positive formal thought disorder, are evident in the answer of a patient (from the current study) to a question about his ruminative thinking:

> *“Ah, I have experienced too much of life, or rather too much of the bad side of life. The ethical questions raised; the Mountain Top speech given on the eve of his assignation by the Rev Martin Luther King, is a topic I reminate [sic] on. The big moral picture; others are entertained by peculiar pictures of cats on the web and what they will eat for tea that evening. I question the means of production of the food I eat; what is the effect on the wild life that shares the farmland; and other tough issues. This is partly why I can feel crushed by the range of cognatation [sic]”*.

Whereas formal thought disorder is clinically defined based on naturalistic speech, loose and atypical associations are also evident in more structured tasks. For example, the above patient produced the following idiosyncratic associations to word cues in a free association task:

Knight –> “*Because”*

Ever –> “*Holiday” T*

Three -> “*Fry”*

Notably, formal thought disorder often involves ‘negative’ symptoms, consisting of a reduction in speech output (Alogia), repetitive speech (Perseveration), concrete thinking, and more. Here we focus primarily on positive thought disorder symptoms (which we refer to as speech disorganization) yet examine specificity by testing associations with negative formal thought disorders symptoms as well.

Neurobiological research on semantic cognition and speech disorganization in schizophrenia has reported structural and functional abnormalities in temporal and frontal brain regions linked to semantic cognition^23–28^. However, such findings offer limited resolution to the longstanding debate about whether semantic structure or process is the core impairment. For instance, abnormalities in brain volume or in structural or functional connectivity may disrupt controlled retrieval processes^24,29,30^, as opposed to causing an impairment in the structural organization of semantic representations. To our knowledge, prior research on the neurobiology of speech disorganization has not examined abnormalities in the *neural encoding of semantic representation*, where the explanatory power of the latter is exemplified in a recent study showing abnormal representation of traumatic memories in PTSD^31^.

To examine how abnormalities in semantic cognition are expressed in schizophrenia, as well as its relations to speech disorganization, we asked how people generate and regulate associations to word cues. Here, atypical associations, as previously reported in schizophrenia^32,33^, could reflect alterations in semantic structure, controlled retrieval processes, or both. Theories of impaired semantic structure in schizophrenia suggest that a participant who reports an atypical association (e.g., the association “Because” to the cue “Knight”; Box 1) might do so on account of these words being closely related^34,35^, or because all possible associations, including more typical ones (e.g., “Armor”) are weakly connected to the cue (‘Atypical’ and ‘Weakened’ structure in Figure 1A)^22^. To probe this aspect of semantic structure, we asked participants to rate the subjective strength of their own associations and leveraged a recently developed method to characterize participants’ personalized semantic structure based on these ratings^36^. We also examined the extent to which semantically related words evoke similar neural representations, as measured by Magnetoencephalography(MEG)^37^. We hypothesized that atypicality in semantic structure would impact an ability to decode neural representations from models of semantic representation trained on population-level corpora^31,38,39^. However, this reduction in decodability should be mitigated when using models that capture individual-level semantic representations. To test the latter, we introduce a novel approach where population-level semantic models are fine-tuned to predict participants’ subjective associative strength ratings and then used for *personalized semantic decoding* analysis.

**Figure 1.**
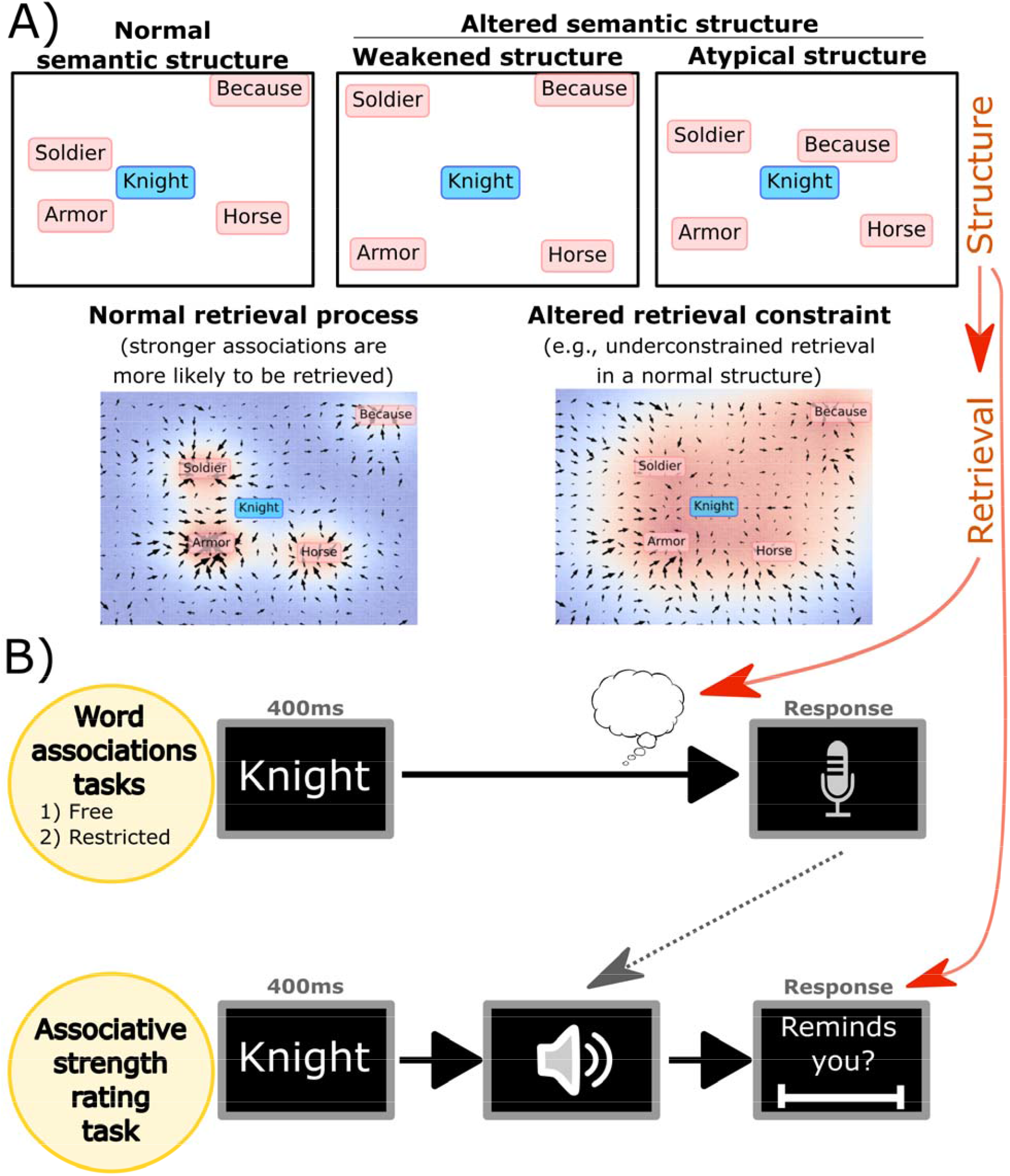
Schematic representation of candidate component processes involved in loosening of association in schizophrenia (Panel A, and red arrows) and the specific tasks we used to capture these processes (B). Panel A depicts a schematic semantic space, and putative retrieval dynamics that may operate on it, illustrating how loose associations can arise out of aberrant semantic structure (particularly reflecting weakened semantic links, or a predominance of atypical semantic links), or aberrant (e.g., more random) retrieval dynamics. The role of retrieval dynamics can also be isolated by manipulating task instructions (B), here corresponding to tasking participants to report strong and common associations (Restricted associations condition) rather than the first associations that comes to mind (Free association condition). Semantic structure is also expected to manifest in the associative strength ratings participants give to their own associations.

Theories highlighting impairments in the retrieval processes operating on normal semantic structure entail that atypical associations are more likely to come to mind or be reported, even if not (subjectively) related to an eliciting cue (Figure 1A, second row). This could reflect greater retrieval noise^40^, suboptimal search strategies^10,41^, or a deficit in flexibly adapting semantic retrieval to specific task context or demands^42^. In either case, we expect patients to report more atypical associations while also rating such associations as relatively weak. To specifically examine patients’ ability to adapt semantic retrieval to task demands, we compared a condition where participants were asked to report the first association coming to mind (*Free Associations Condition*) with a condition where they were encouraged to report only strong and common associations (*Restricted Associations Condition;* Figure 1B).

## Results

### Schizophrenia patients produce more atypical associations

Thirty-one patients with schizophrenia and thirty-one controls (see Table S1 for clinical and demographic information) completed a word association task during simultaneous acquisition of MEG data. On each trial, participants were presented with a cue word and asked to generate either Free, or Restricted (i.e., strong and normative) associations. In a healthy control group, the latter instructions successfully reduced production of atypical associations (95% odds ratio Bayesian credible intervals = [0.80, 0.91]; Figure 2A). Here, atypical associations correspond to associations that no other participant (in previous norms or our study) reported in response to that cue. We focused on this norms-based measure, rather than similarity metrics extracted from language models (cf.^41,43,44^), as the latter was less sensitive to our key manipulation (Cohen’s *d*’s = 0.68 and 0.38, respectively; but see Supplementary Note S1 for analyses using this alternative measure)

**Figure 2.**
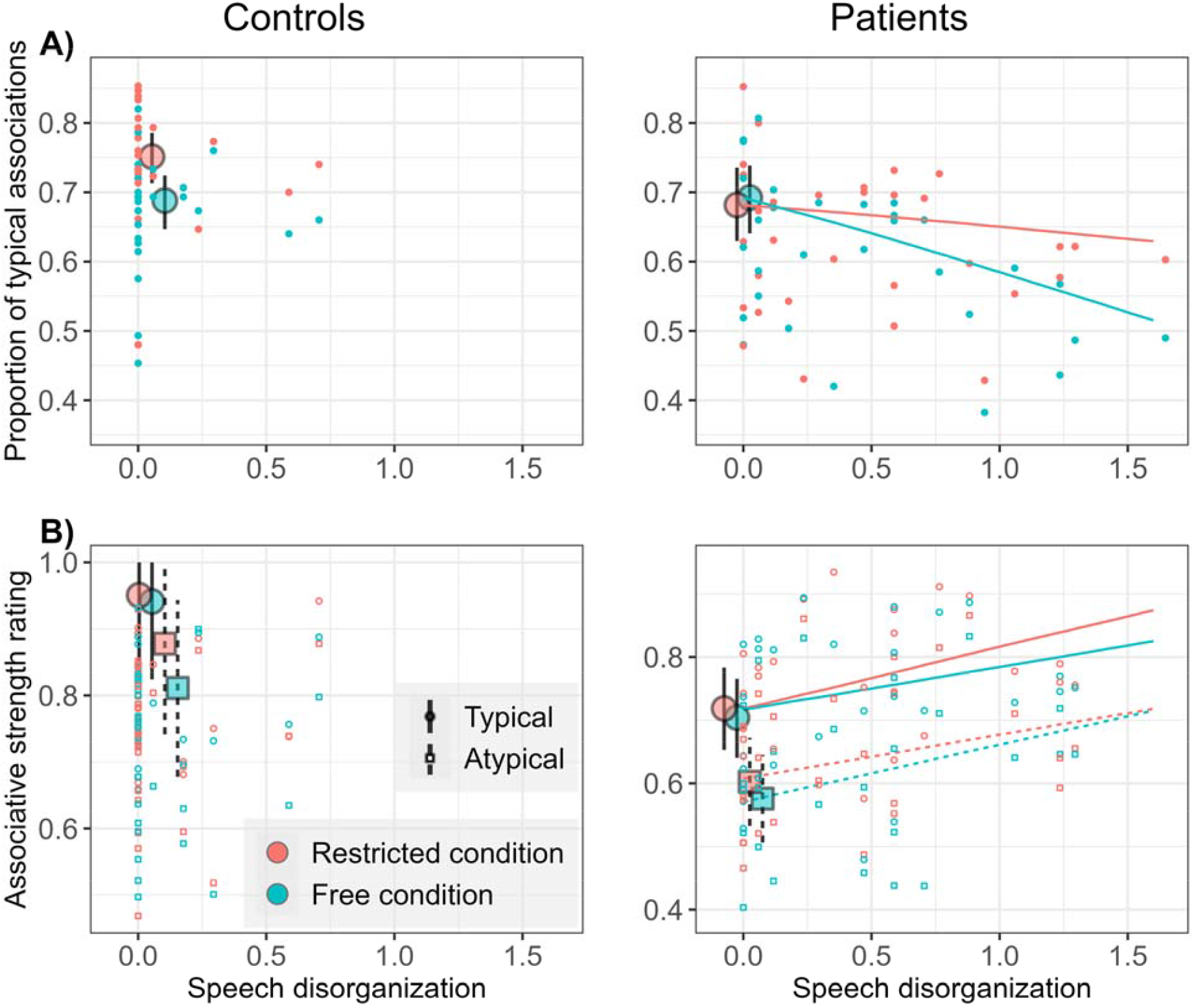
Effects of schizophrenia diagnosis (colored boxplots) and speech disorganization (X-axis) on the objective and subjective strength of emitted associations, measured based on their typicality and subjective ratings, respectively. Patients with speech disorganization symptoms produced more atypical associations in the Free condition, but rated their associations as similarly to controls. Conversely, patients with minimal speech disorganization mainly exhibited a difficulty restricting associations when required to do so, while rating their associations as weak, in comparison to controls.

We determined the degree of speech disorganization, independently from the task, based on a clinical interview using the Thought and Language Disorder scale^45^. Additionally, the specific role of speech disorganization (also known as positive formal thought disorder) was determined by contrasting it with the effects of a pathological reduction in speech output (i.e., negative formal thought disorder), as well as the effects of general symptom severity measured using the Positive and Negative Syndrome scale^46^ total score. Finally, to separate the effects of diagnosis from those of speech disorganization, the former was examined while statistically fixing speech disorganization to zero.

Both diagnosis and speech disorganization symptoms predicted greater atypicality in associations, but these factors showed distinct patterns depending on task condition (Fig. 2A). In particular, patients (with low speech disorganization symptoms, *SDis*) showed a reduced ability to minimize atypical responses in the Restricted condition (*OR*_*groupXcondition*|*Sdis=0*_ = [1.01, 1.11], *OR*_*condition*|*patients,Sdis=0*_ = [0.90, 1.02], *OR*_*condition*|*controls,Sdis=0*_ = [0.80, 0.91]), but displayed no increase in atypicality in the Free condition (*OR*_*group*|*Sdis=0, free*_ *=* [0.88, 1.13]). Conversely, speech disorganization symptoms (SDis) predicted greater atypicality in the Free condition, but not in the Restricted condition (*OR*_*SDisXcondition*|*patients*_ = [0.74, 0.98], *OR*_*Sdis*|*free*_ = [1.14, 2.13]; *OR*_*Sdis*|*restricted*_ = [0.78, 1.56]). Testing specificity with regards to other symptoms indicated that neither a reduction in speech output (negative thought disorder), nor general symptom severity showed a clear correlation with association atypicality (*OR* = [0.98, 1.35], *OR* = [0.99, 1.02], respectively), nor an interaction with condition (*OR* = [0.93, 1.11], *OR* = [0.99, 1.01]).

Together, these results indicate that, in schizophrenia, basic atypicality in word associations is a specific marker for speech disorganization, where this relationship cannot be reduced to a difficulty in adapting speech to goals or task instructions. This contrasts with the effect of diagnosis itself, particularly for patients with minimal speech disorganization, who exhibit a specific difficulty in restricting associative output when required to do so.

### Distinct alterations in semantic structure in patients with and without speech disorganization

We next investigated individual differences in the semantic structure underlying associations. For this purpose, after the word association task participants were tasked to rate the strength of their own generated associations (henceforth: *Associative Strength ratings*). A crucial prerequisite in comparing patients’ ratings to those of controls is to ensure that patients’ ratings are not excessively biased or noisy. We examined the former by collecting ratings for random pairings as well as, in a separate task, probing similarity between colors (see Methods). The latter was examined by collecting repeated ratings for a subset of cue-association pairs. Patients did not differ from controls in ratings of random pairings (CI_group_= [−0.12, 0.10]), or independent colors (CI_group_= [−0.02, 0.08]), providing no evidence of bias. Patients’ ratings were no less reliable than those of controls’ 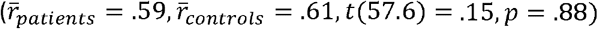. Note, however, that since four controls and three patients showed negligible reliability (r < .25) they were removed from analyses related to associative strength.

Under the hypothesis that patients’ atypical associations reflect primarily aberrant retrieval processes operating over a normal semantic structure, we would expect such atypical associations to obtain low associative strength ratings. Conversely, atypical associations could also reflect an atypical semantic structure (Figure 1A), in which case we would not expect patients to experience such associations as weak. Finally, a general weakening of associative links (Figure 1A) should be reflected in a global reduction of associative strength, evident for both typical and atypical associations. In accordance with the latter, patients tended to perceive their associations as relatively weak (CI_group|SDis=0_= [−0.20, −0.04]), regardless of condition (CI_groupXcondition|SDis=0_= [−0.02, 0.01]), or association typicality (CI_groupXtypicality|SDis=0_ = [−0.03, 0.02]). Conversely, speech disorganization symptoms did not significantly correlate with associative strength ratings, and instead they showed a positive trend (CI_SDi’s_= [−0.05, 0.21]). Thus, although patients with speech disorganization produce more atypical associations (Figure 2A) they do not experience these (or other) associations as exceptionally weak (Figure 2B), in keeping with the hypothesis of atypicality in semantic structure that goes beyond a general weakening.

To further probe abnormalities in semantic structure, we leveraged a recently developed method for inferring individuals’ associative structure through generative modeling^36^. This approach assumes that an individual’s associative structure includes both typical associations (derived from norms), and idiosyncratic (i.e., atypical) associations which differ between participants and cues. Intuitively, if associative retrieval is conceptualized as random sampling from memory, then the number of *possible* atypical associations (for a given participant and cue) can be probabilistically inferred based on the number of *reported* atypical associations. However, atypical associations are usually weaker (Figure 2B) and their expression can be regulated depending on context (i.e., in the Restricted condition). Thus, inferring the number of possible atypical associations from reported atypical associations requires partialing out these effects to avoid underestimation (Figure 3A). In practice, this procedure involves availing of a set of generative Bayesian regression models to estimate both the number of atypical associations for each cue and participant and their respective strengths (see *Methods*). The validity of this method has been established in previous studies^36,47^, and again here, using model recovery simulations (Methods). As further validation, we found that the estimated number of atypical associations predicts the degree of competition among associations, as reflected in RTs, even controlling for the number of typical associations and the typicality of the reported association (CI = [0.001, 0.053]). Finally, as expected, this procedure revealed reported associations consist of the upper tail of a distribution of the associations participants could have reported (Fig. 3B).

**Figure 3.**
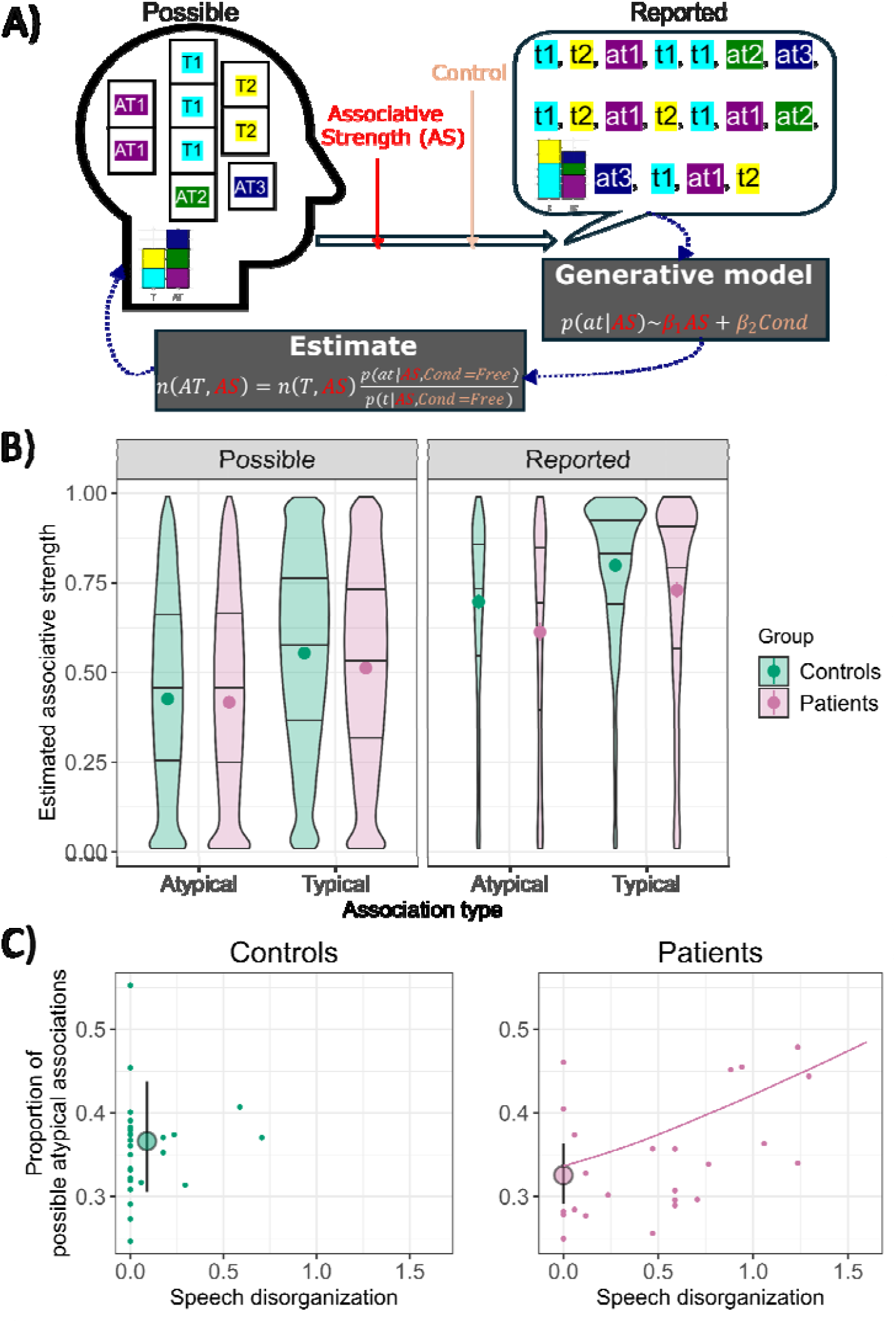
Estimated associative structure. Panel A illustrates the estimation method, by depicting an urn model of possible associations (for a given cue and participant). Here, possible associations include two typical (T1, T2), and three atypical associations (AT1, AT2, AT3), wherein associative strength is illustrated by the number of ‘balls/colors’ representing each association. Reported associations reflect one possible sample of associations, wherein stronger associations are more likely to be sampled. Whereas the overall proportion of reported atypical associations (p(at)) does not generally reflect the proportion of possible atypical associations (n(AT)), because atypical associations are weaker and their expression is regulated (in the Restricted condition), we can partial out these effects using generative modeling. Panel B depicts the distributions of the estimated strength of possible (and, for comparison, reported) associations (the area reflects the respective number of possible and reported associations). Horizontal lines and points represent quartiles and means, respectively. Panel C depicts the estimated number of atypical associations in participants’ associative structures as a function of diagnosis and speech disorganization symptoms.

Analyzing participants’ estimated associative structures provided additional support for a general weakening of associative links in patients (without speech disorganization) in comparison to controls, as reflected in the average estimated strength of possible associations (CI_group|SDis=0_= [−0.05, −0.01]; Figure 3B). Conversely, speech disorganization in patients correlated with a greater proportion of possible atypical associations (CI_SDis_ = [0.06, 0.64]; Figure 3C). We did not observe similar atypicality when comparing patients with minimal speech disorganization to controls (CI_group|SDis=0_ =[−0.29, 0.10]). In support of specificity, neither a reduction in speech output among patients (negative thought disorder), nor general symptom severity were associated with estimated atypicality of associative (Cis = [−0.23, 0.20], [−0.01, 0.02], respectively).

Overall, these results suggest that diagnosis and speech disorganization symptoms relate to distinct abnormalities in semantic structure. The atypical semantic structure in patients with speech disorganization can also explain why their tendency to report atypical associations is more evident in the Free condition, where controlled retrieval dynamics play a lesser role. Conversely, while diagnosis (when controlling for speech disorganization symptoms) related to a general weakening of associative links, it remains to be determined whether this is associated with a difficulty in regulating typicality of retrieved associations found in the same individuals.

### Less constrained retrieval in schizophrenia

As we have shown, patients with low speech disorganization are characterized by a general weakening of associative links, and this weakening was not moderated by condition or associative typicality. On the other hand, these same patients exhibited a specific difficulty adapting their reported associations to task demands, suggesting an additional impairment at the level of controlled retrieval. To identify specific impairment of processes constraining retrieval, we examined whether the associations that patients reported were both weaker and more atypical than what would be expected based on their estimated associative structure. To examine the strength of reported associations, we computed the *relative strength* of reported associations, defined here as the percentile rank of the reported association within the individual’s estimated space of possible associations to a given cue. Conversely, to examine their typicality, we computed the *relative atypicality* of reported associations, defined as the extent to which a participant is more likely to report an atypical association on a given trial than what is expected based on the probability mass of atypical associations for a given participant and cue.

Prior to analyzing individual differences, we first examined whether relative strength and relative typicality increase in the Restricted condition, when participants are tasked to downregulate weak and atypical associations. As expected, the relative strength of controls’ associations increased in the Restricted condition (CI_condition|controls_= [0.02, 0.09]), whereas relative atypicality decreased (CI_condition|controls_= [−0.23, −0.09]). Consistent with an alteration in the retrieval process that goes beyond structural abnormalities, relative strength was lower among patients (CI_group|SDis=0_ = [−0.21, −0.02]), regardless of condition (CI_groupXcondition|SDis=0_ = [−0.04, 0.01]). Similarly, relative atypicality was higher among these patients (CI_group|SDis=0_ = [0.02, 0.14]), although we also found an interaction (CI_groupXcondition|SDis=0_ = [−0.01, 0.12]) indicating that patients (in contrast to controls) failed to sample more typical associations in the Restricted condition, even when controlling for structural typicality (CI_condition|patients, SDis=0_ = [−0.10, 0.04]). Notably, speech disorganization symptoms among patients did not correlate with relative strength (CI_SDis_= [−0.33, 0.20]), or relative atypicality (CI_SDis_= [−0.16, 0.17]).

### Atypical semantic structure in speech disorganization is reflected in neural representation

The above findings converge on an interpretation that speech disorganization in schizophrenia is associated with an underlying atypicality in semantic structure. To further examine abnormalities in semantic structure, independent of participants’ behavioral responses, we examined the neural representation of cues (to which participants produced associations). While semantically related words tend to evoke similar neural responses^3,37,39,48^, atypicality of semantic structure can be expected to corrupt this similarity mapping. To formally test this, we used a similarity-decoding framework^37^, which leverages the benefits of representation similarity analysis (RSA; e.g., no tunable parameters, comparing different semantic models) while producing a cross-validated estimate of decoding accuracy.

For this analysis, the neural representation of a specific cue is defined by reference to the correlation between the MEG response it evokes, and the responses evoked by all other cues. Likewise, the semantic representation of a cue is defined by measuring its similarity to other cues using semantic models trained on large text corpora. It follows that the decodability of a cue can be indexed by the extent to which its neural representation correlates to its semantic representation than to the semantic representation of every other cue, using a leave-2-out cross-validation (Figure 4a)^37^.

**Figure 4.**
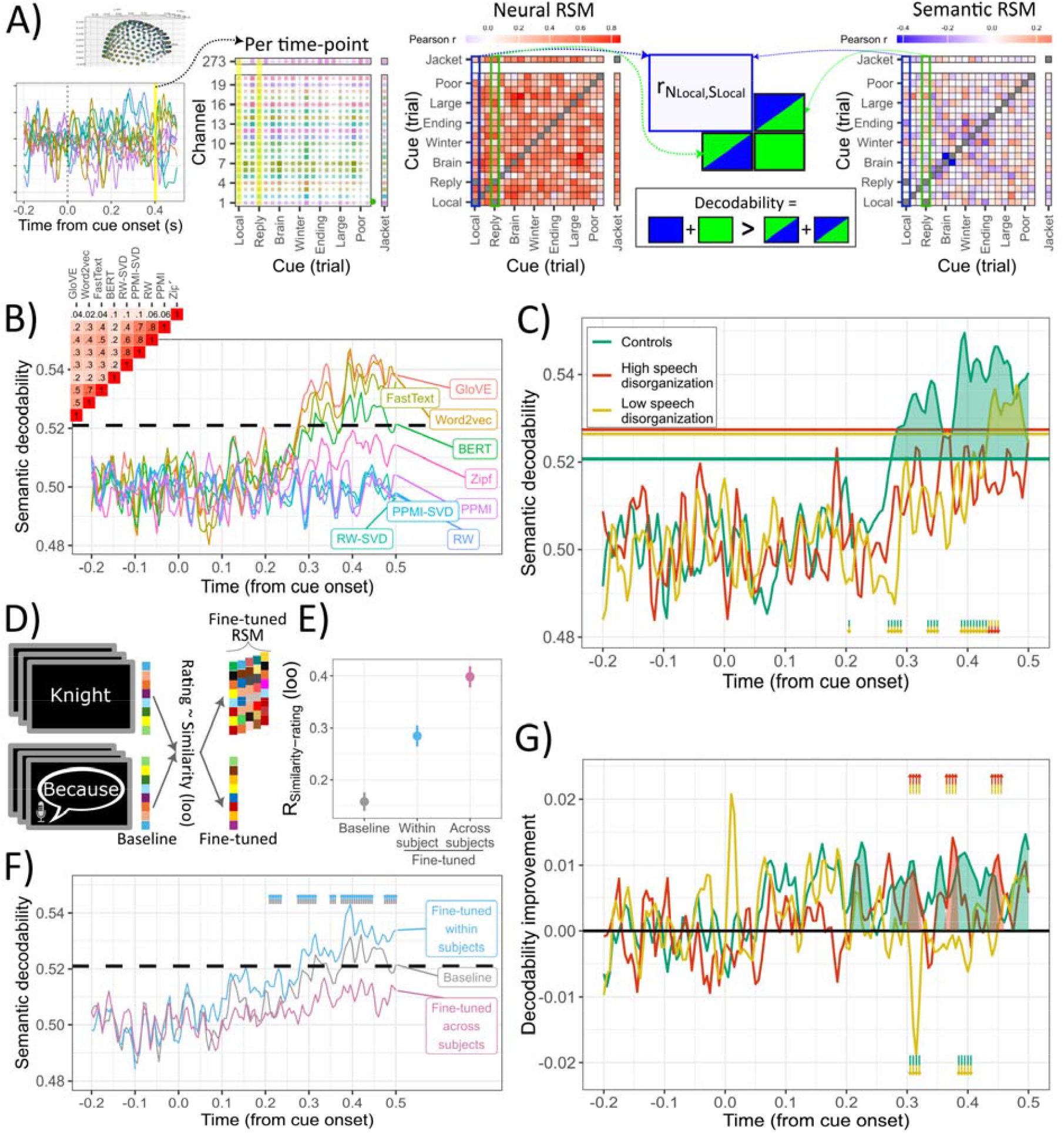
Method and results of a semantic-decoding analysis. Neural representational similarity matrices (RSM) were calculated as the Pearson correlation in spatial patterns among cues for each timepoint (A). Semantic RSMs were based on several semantic space models trained on large text corpora (GloVE, Word2vec, FastText, DistilBERT), or free association norms (RW-SVD, PPMI-SVD, RW, PPMI; Panel B). As a control analysis, we ensured semantic decodability was not explained by simple word frequency effects (Zipf; Panel C). Following model-specific decodability results (B), group differences (Panel C) were examined by averaging RSMs across GloVE, Word2vec, FastText. Shaded areas represent above-chance decodability. Arrows represent significant differences between groups, while controlling for speech disorganization (green to orange arrow), or a significant effect of speech disorganization (orange to red arrows), detected using a cluster permutation test. To approximate subject-specific semantic models, DistilBERT embeddings were fine-tuned to predict subjective associative strength ratings, using a leave-one-out procedure (Panel D). Fine-tuning within and between-subjects improved the alignment between model embeddings and ratings (Panel E). Conversely, only within-subject fine-tuning improved semantic decodability in controls (Panel F). Analyzing individual differences in the latter indicated that semantic decodability improved only for patients with speech disorganization symptoms (Panel G). RW – random walk; PPMI – positive pointwise mutual information; SVD – singular value decomposition; Loo – Leave-one-out

Neural representational similarity codes (RSM) were calculated using a temporal searchlight approach, correlating spatial patterns across cues (Figure 4a). We focused on the 500ms following cue onset to ensure we captured semantic processing of the cue^49,50^, thereby minimizing processes that relates to a subsequent retrieval of associations. Semantic similarity was calculated using several common semantic models, based on either large text corpora (e.g., GloVE, Word2vec, FastText, DistilBERT) or measures derived from large free association datasets^51,52^. Whereas different semantic models correlated with one another (Figure 4B), only models based on text corpora predicted neural representations (Figure 4B), with no clear differences between the three top models (GloVE, Word2vec, FastText). On this basis, we opted to use an aggregated measure of semantic similarity by averaging across these models.

We found above-chance semantic decodability among healthy controls, beginning ∼280ms after cue onset (Figure 4C). More importantly, as hypothesized, speech disorganization symptoms among patients were associated with reduced semantic decodability (orange-red arrows and red line in Figure 4C). Conversely, neither a reduction in speech output (negative thought disorder), nor general symptom severity related to decodability (Figures S2-S3). Note, however, that a reduction in semantic decodability was evident even among patients with low speech disorganization (turquoise-orange arrows and orange line in Figure 4C). This unexpected finding may indicate a degree of atypicality in neural encoding of semantic representations even among patients without speech disorganization. Alternatively, this reduction in semantic decodability might reflect other causes affecting the reliability of the neural signal. In keeping with the latter interpretation, controlling for head movement during recording, and to a lesser extent, other estimates of neural signal-to-noise ratio, mitigated the reduction in decodability seen in patients with low speech disorganization (but not the effects of speech disorganization; Figure S1). Thus, although we found alterations in the neural encoding of semantic representations in both patients with and without speech disorganization, the latter was partially explained by a reduced reliability of the neural signal rather than an atypicality of semantic structure per se.

To further examine the specific role of atypical semantic structure in reducing semantic decodability, we next examined decodability using personalized semantic models. Thus, a reduction in decodability explained by atypical semantic structure should be mitigated when using personalized rather than population-level semantic models. To examine this hypothesis, we used DistilBERT, which (like other transformer models) not only allows quantifying similarity between words (or sentences) but can also be fine-tuned without extensive training data^53,54^. We fine-tuned the model, separately for each participant, such that similarity between the embeddings of a cue and an association match participants’ subjective associative strength ratings, using a leave-one-out procedure to avoid over-fitting (see Figure 4D and Methods).

As depicted in Figure 4E, fine-tuning successfully modified the embeddings of the relevant cues and associations to increase the correlation between similarity and subjective ratings. Furthermore, using the modified cue embeddings increased semantic decodability among controls, relative to a generic (baseline) DistilBERT model (gold-turquoise arrows in Figure 4F). Crucially, this improvement in decodability is best explained by the procedure’s success in approximating personalized semantic models, rather than capturing general task features, as indicated by a lack of improvement in decodability when fine-tuning the model across participants. Finally, as hypothesized, using the fine-tuned (vs. baseline) model improved decodability among patients with speech disorganization, while doing the opposite in patients with minimal speech disorganization. This finding, that personalized semantic models improve decodability in those with speech disorganization, supports the hypothesis that reduced decodability in speech disorganization is, at least partially, explained by atypicality of semantic structure. That said, fine-tuning did not bring decodability in those with speech disorganization to normal levels (which would require greater improvement in decodability among those with speech disorganization versus controls), suggesting additional factors that may contribute to reduced decodability in schizophrenia in general.

## Discussion

A key challenge in the study of semantic cognition, thought, and language is the intricate interplay between underlying conceptual structure and retrieval processes acting on this structure. Indeed, we conventionally measure structure by invoking a downstream retrieval process, leading to an interpretational ambiguity^1,4,7,13^. Such complexities are accentuated in schizophrenia, where disorganized speech, known as positive formal thought disorder, is prevalent; yet, the mechanistic relationship between the latter and structural alterations in conceptual representations on the one hand, and retrieval processes operating on these structures on the other, has remained elusive.

In this study we provide converging behavioral and neural evidence consistent with an hypothesis that speech disorganization symptoms in schizophrenia relate to atypicality in semantic structure. Behaviorally, patients with speech disorganization produce more atypical associations and show less insight concerning the atypicality of their associations. Furthermore, a finding that speech disorganization was associated with greater mismatch between neural representation and population-level semantic models, but that this reduction was partially mitigated when using personalized, semantic models provides convergent support for an account based on atypicality of semantic structure.

Schizophrenia was associated with alterations in semantic cognition even after controlling for speech disorganization symptoms, but notably this appears to involve a distinct mechanism. At the absence of speech disorganization, patients were characterized by more disconnected and weakened semantic structures, as well as less constrained retrieval processes. The latter was particularly evident in their specific difficulty adapting associative output to task demands. Whereas these individuals also exhibited alterations in neural representations, these were better explained by neural noise (particularly head movements) than reflective of atypicality in semantic structure.

Previous attempts to dissociate semantic structure and process in schizophrenia deployed tasks probing semantic knowledge (e.g., asking to define or categorize ‘Knight’), where impairments in retrieval (but not structure) might be assumed to engender inconsistent performance^16,17,22^. However, such methods, originally developed for neurological conditions where semantic degradation is common (e.g., semantic dementia)^2,12,20,55^, are arguably less suitable for schizophrenia research. Indeed, people with schizophrenia can manifest marked disorganization and incoherence in discourse, with no reduction in vocabulary or simple comprehension^56^. Therefore, it is not surprising that previous attempts using such methods found inconsistent results^16,17,22^. Here, we detail an alternative approach, one that enables an analysis of the interplay of structure and retrieval in a spontaneous language production task, which bears greater similarity to natural discourse. Thus, rather than arbitrating between structural and controlled retrieval impairments, this enables an examination of how the two types of impairments interact.

From a neural perspective, previous work on schizophrenia has revealed abnormalities in the structure and function of semantic brain areas^23,24,27,57^, as well as alterations in electrophysiological markers of semantic processing^58–60^. While such univariate markers of semantic processing greatly contribute to our understanding of semantic cognition in schizophrenia, they do not capture how semantic information is *represented* in the brain per se. Here, we go beyond merely identifying abnormalities in semantic processing to show how these abnormalities reflect reduced alignment with normative semantic structures. To determine the extent to which this misalignment reflects atypicality in semantic structure, we developed a novel approach where neural representation is mapped to personalized (potentially atypical), rather than normative, semantic structure. Establishing a method for decoding neural representations at an individual level has potential to advance a nascent line of research that seeks to explain mechanisms of psychopathology in terms of abnormalities in neural representation^31^.

Our findings highlight two important theoretical questions. First, is there a causal relationship between weakening of associative links and a reduced constraint over associative retrieval seen in patients? Second, does the emerging dissociation between the effects of diagnosis and the effects of speech disorganization symptoms reflect stable subtypes, or rather dynamic adaptation processes whereby atypical semantic structure arises through compensation for an initial weakening of semantic structure? Placing our findings within the framework of previous neurocomputational models of schizophrenia can aid in formulating well-founded, albeit speculative, answers to these questions.

Our finding of weakened associative links resonates with seminal work on the disintegration of cognitive functions^61^ and neural dysconnectivity in schizophrenia^27,62^. Intriguingly, more recent computational characterizations of neural dysconnectivity in schizophrenia highlight implications for reduced stability of attractor networks^40,63,64^, or, from a Bayesian perspective, a less precise and less stable model of the world (i.e., reduced ‘prior precision’^65–67^). These, in turn, could lead to a greater influence of external input and internal noise^40,63,64^, and greater behavioral stochasticity^68,69^. Thus, greater retrieval stochasticity may not simply co-occur with a weakening of associative links among patients without speech disorganization but instead reflect a natural consequence of attractor networks characterized by non-linear rich-get-richer dynamics^36^.

In this context, our finding that speech disorganization can be explained by (potentially stable) structural atypicality is surprisingly similar to a previously noted contradiction between attractor instability and inflexible false beliefs (i.e., delusions) in schizophrenia^70^, formalized in Bayesian models as a tension between reduced versus increased prior precision^66,71,72^. Recent work has suggested, however, that delusions can arise in an imprecise cognitive system due in part to standard learning mechanisms, implemented over time^71,73–75^. Along similar lines, the emergence of atypical semantic structure in patients with speech disorganization could arise out of internal learning dynamics, whereby retrieving weak, under-constrained associations may eventually strengthen them at the expense of more typical associations^76,77^. This may also shed light on the results of two recent studies suggesting that semantic predictions are less stable, yet act as stronger priors for auditory processing in schizotypy^78,79^. From a neurobiological perspective, this speculative explanation aligns with the idea that an initial loss of synaptic plasticity in schizophrenia (due to NMDA receptor dysfunction^80–82^) leads to a compensatory reduction in neural inhibition^83–85^which, in-turn increases synaptic plasticity at the cost of strengthening more random connections.

The distinction between structural and process-related impairments in semantic memory has important clinical implications. An atypical structure could be less flexible and less amenable to change, potentially explaining why speech disorganization in schizophrenia has been associated with greater difficulties in social functioning, lower insight, and reduced treatment prospects^19,23^. Here, the idea that observable disorganization in discourse could be mechanistically preceded by more subtle alterations in semantic cognition has important implications for recent suggestions that language can serve as a biomarker for psychosis^33,86–88^. Thus, by measuring subtle alterations in semantic cognition, even if discourse disorganization is (still) lacking, researchers might improve the timely delivery of preventive interventions.

We acknowledge several limitations to our study. First, formal thought disorder is a multidimensional construct, involving linguistic, cognitive, and social components^23,77,89^. Here we focused on single-word associations, while predominant accounts of formal thought disorder focus on impairments in integrating a higher-level context (e.g., at the level of sentences) in language production and comprehension^77,86^. Second, our analysis of the neural markers of semantic structure in schizophrenia and formal thought disorder did not focus on identifying *where* or *how* semantic information is represented in the brain – questions extensively addressed in past studies^2,23,90^. Instead, relying on a common assumption that semantic neural activation is multimodal and globally distributed^2^, we sought to identify individual differences in *what* semantic information is represented in the brain.

Overall, our study demonstrates the power of a theory-driven, multi-method approach in characterizing mechanisms of semantic cognition and its disturbances. By revealing that speech disorganization in schizophrenia arises from abnormal semantic structure, while more intact speech may mask underlying deficits in controlled retrieval processes, we reveal a dissociation with significant theoretical and clinical implications. Although grounded in schizophrenia, this framework offers a generalizable approach for probing how semantic cognition breaks down across a range of disorders^2,20^ and how it also deviates in non-pathological contexts, such as creativity^4,13^. Furthermore, the proposed personalized semantic decoding could greatly benefit the rapidly growing literature using language models to understand linguistic representation in the brain^37–39,91,92^. Here, an ability to fine-tune modern language models to specific individuals without extensive data may increase decoding accuracy, as well as help dissociate the common and unique components of linguistic representations. From a more theoretical perspective, our findings suggest a possible bi-directional relationship between structure and process, where patterns of retrieval not only reflect semantic structure but may also reshape it over time. Understanding how these dynamics unfold, especially in social communicative contexts, remains a pressing frontier for research on both typical and atypical semantic cognition.

## Methods

### Participants and assessment

The study was approved by the London Westminster NHS Research Ethics Committee (15/LO/1361). All participants provided written informed consent and were compensated for their time. We recruited 33 patients with schizophrenia (n = 25) or schizoaffective disorder (n = 8) from London and Surrey psychosis NHS clinics, and 32 healthy controls from the same geographical area through online advertisements. General exclusion criteria included comorbid neurological disorders (including autistic spectrum disorders), current suicidal intent, or cognitive or motor difficulties prohibiting the patient from remaining seated and focused for 2 hours during the MEG scan. Healthy volunteers were not taking neurological or psychiatric medication, had no history of neurological or psychiatric disorder, and no family history of psychosis. The final sample comprised 31 patients (two patients dropped out before the word association task), and 31 controls (one control was unable to keep awake during the task).

All participants underwent a semi-structured psychiatric interview (MINI-V^93^), as well as a structured assessment of depression (Montgomery Åsberg Depression Rating Scale^94^). Verbal IQ was assessed using the Weschler Test for Adult Reading (WTAR^95^). Patients’ psychiatric symptoms were additionally assessed with the Positive and Negative Syndrome Scale (PANSS^46^). Finally, formal thought disorder symptoms were assessed using the Thought and Language Disorder scale (TALD^23^). Cognitive and clinical assessments were conducted prior to MEG scan.

Groups were matched on age, gender, education, ethnicity, and country of primary education (Table S1). Whereas patients had a lower verbal IQ (95% CI = [2.17, 15.18]), these differences were statistically controlled for in the above analyses. Twenty-nine patients were medicated, whereas ten patients had comorbid disorders (primarily Panic Disorder and Obsessive Compulsive Disorder; Table S1). See Table S1 for full demographic, cognitive and clinical scores. All participants were asked to undergo a mouth swab drug test, and all 56 participants who agreed showed no trace for recent substance use.

### Experimental procedure

#### Word association task

Participants were presented with cue words and asked to vocally report an association to that cue. The task consisted of two conditions. In the Free Associations Condition, participants were asked to report the first association that came to mind. In the Restricted condition participants were asked to report only strong and common associations. The two conditions were presented in fixed order to avoid carry over effects (from Restricted to Free). Each condition consisted of 150 trials (with short breaks after 60 and 120 trials), and both preceded by training trials (15 and 20, respectively). Participants responses in the latter condition were examined in real-time and considered correct if they were consistent in previous free association norms^52,96^ whereas percentage accuracy feedback was presented every twenty trials.

Each trial started with a fixation cross, presented for 750-1000ms, followed by a cue word (e.g., table) presented at the center of the screen for 300ms. Participants were asked to say out loud an association and press the ENTER key once finished. Reaction times were determined offline based on vocal onset using Matlab’s *detectSpeech* function, which uses a thresholding algorithm based on energy and spectral spread per analysis frame. The Word association task was followed by a filler task wherein participants saw pairs of related or unrelated words^97^ followed by a break. After this, participants completed a Rating task designed to estimate the subjective associative strength of their reported associations. Thus, on each trial participants were presented with a cue that was followed by auditory playback of the association they produced in response to that cue during the Word association task. Cues were presented in random order to minimize participants’ ability to remember when (e.g., in which condition) that cue was presented. In addition, to ensure participants did not rely on their own vocal onset, thinking time was programmatically removed from the recordings participants listened to. For each cue-association pair, participants were asked to determine the extent to which the cue word reminds them of the association, based on their subjective intuition and knowledge (e.g., ignoring what other people might think). Associative strength (AS) ratings were collected using a continuous visual analog scale from 0 (“not at all”) to 1 (“very much”).

To measure reliability of the rating task, we repeated 20 trials, and measured the Pearson correlation between the two ratings. In addition, the associations played on 20 trials were shuffled (i.e., came from a different cue), allowing us to both measure general biases in ratings and detect random responding (where the distribution of ratings on such shuffled trials is expected to overlap with that of normal trials). These quality control measures led us to exclude 7 participants (3 patients) from analyses involving the associative strength ratings, in addition to two participants (1 patient) who decided to drop out prior to the ratings phase.

#### Color similarity task

The color similarity task was designed to measure general biases in rating similarity. On each trial, participants were presented with two colors and asked to rate their similarity, with instructions that mirrored those used in the associative strength ratings task. The task included 48 trials including different combinations of blue and green shades. The task was completed by 54 participants (28 patients).

### Analytical procedures

#### Statistical analyses

Generalized mixed effects models were used for most statistical analysis. Our strategy for distinguishing the effects of schizophrenia diagnosis from the specific effects of speech disorganization symptoms (found primarily in the schizophrenia group) involved running two separate models for each dependent variable. Thus, the specific effects of speech disorganization were examined within the schizophrenia group. Conversely, the effects of diagnosis, separated from speech disorganization, was tested by controlling for speech disorganization symptoms (while fixing speech disorganization in the control group to zero), such that the group effect can be interpreted as the predicted difference between patients with minimal speech disorganization and controls. Generalized mixed effects models were fit using Hierarchical Bayesian framework (R brms package^98^) based on a number of considerations. First, associative strength ratings were parameterized using a rectified normal distribution, reflecting an assumption that participants may sometimes prefer to give ratings above 1 (or below zero) but are unable to do so. Implementing this parameterization is not feasible using maximum likelihood approaches. Second, Bayesian hierarchical models usually produce more stable^99^ estimates in complex mixed effects models, and tend to be more robust to multicollinearity^100,101^. Analysis focused on typicality used a logistic regression. Analysis focused on the proportion of atypical associations used a Beta regression, with a log link.

#### Estimating individualized spaces of associations

We estimated the space of possible associations for each cue and participant by combining information from previous word association norms, with subjective associative strength ratings, and the empirical proportion of atypical associations. For a given cue, let *n(T)* and *n(AT)* denote the number of possible typical and atypical associations. We calculated the number of possible typical association (*n(T)*) from the distribution of associations reported for that cue in both previous norms^52,96^, and our study (by at least two participants, to exclude atypical associations).

The probability of reporting an atypical association (*p(at)*), depends on the number of possible atypical associations and their relative associative strengths (AS). Thus, since atypical associations tend to be weaker than typical associations^36^, the proportion of reported atypical associations tend to underestimate the proportion of possible atypical associations. This also means that conditioned on associative strength, the proportion of reported atypical associations can be used to estimate the number of possible atypical associations:

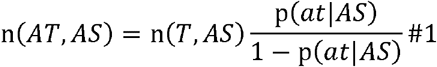

Because associative strength is continuous, we model the odds of reporting a non-normed association as a function of associative strength rating using mixed-effects logistic regression. Using this approach obviates the need to collect multiple associations with equal associative strength for each cue and participant, and yet, can account for differences between participants and between cues in the odds of reporting a non-normed association:

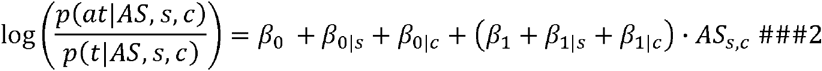

Next, to estimate the associative strength for unreported typical associations, we rely on the relationship between associative strength ratings and the proportion of participants reporting each normed association in previous norms (*pTP*). Thus, we use a linear mixed-effect model to predict the expected AS of association of participant *s* in response to cue *c* as:

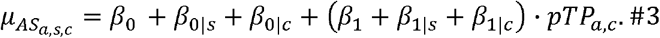

To account for the limited scale of AS ratings [0,1], we assume it is sampled from a rectified normal distribution. Note that Equations 2 and 3 included additional fixed effects for group and condition, and all possible interactions (which are not presented here for brevity).

Both mixed-effects models (Equations 2-3) were estimated jointly in a hierarchical Bayesian model, using Hamiltonian MCMC as implemented in Stan^101^, with non-central parameterization, and weakly information priors. The MCMC included three chains, with 3,000 iterations each, 500 of which were discarded during warm-up to calibrate the Hamiltonian parameters. No divergent transitions were detected, indicating unbiased sampling. Similarly, all models converged as indicated by 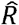 values the did not exceed 1.01.

The above procedure allows us to estimate the associative strength of each possible typical association and the odds that an association is atypical for each possible AS. Using these, we can approximate the number and strength of atypical associations. To do this, we split the range of AS ([0,1]) into small bins (of size 0.02; using smaller bins does not alter the results^36^) and then estimate via sampling the expected number of typical associations within each bin, denoted by 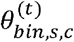. Crucially, since condition is expected to modulate the proportion of reported atypical associations, this effect was estimated in the Bayesian mixed-effects regressions delineated above but removed when estimating the number of possible atypical associations within each bin. Then, the estimated number of atypical associations within each bin is given by:

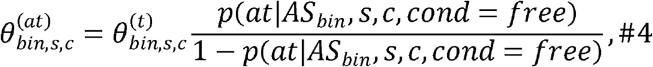

The resulting (non-integer) expected numbers of possible associations for each AS bin is then used in a simple discretization procedure to derive a discrete space of possible associations^36^.

Evidence for the recovery and validity of this method has been established in previous studies^36,47^, and in the current data. Here, we tested recoverability by randomly sampling novel associations from the estimated structure and applying the delineated method to the sampled associations. Associations were sampled in proportion to their associative strength, and a constant weighting factor reducing the sampling of atypical associations in the Restricted condition to match the empirical results. This analysis showed a strong correlation between estimated and recovered number of non-normed associations in both conditions (r_Free_ = 0.89; r_Restricted_ = 0.89), as well as strong correlations between the estimated and recovered values of the average associative strength of possible atypical associations (r_Free_ = 0.74; r_Restricted_ = 0.74).

#### MEG acquisition and preprocessing

MEG was recorded continuously at 600 samples per second using a whole-head 275-channel axial gradiometer system (CTF Omega, VSM MedTech), while participants sat upright (2 sensors not recorded due to excessive noise in routine testing). Three coils were used to locate the position of participant’s head in the three-dimensional space with respect to the MEG sensor array. These were placed on the three fiducial points: the nasion and left and right pre-auricular areas. The coils generate a small magnetic field used to localize the head and enable continuous movement tracking. An Eyelink eye-tracking system was used to monitor participant’s eye movements and blinks. The task was projected at 60Hz onto a screen suspended in front of the participants.

Preprocessing was conducted using MNE-python.^102^ First, head movements were measured and then partially corrected using Maxwell filtering^103^. Next, sensor data were bandpass filtered (FIR filter using the window method) from 0.01 to 40Hz to remove slow-drifts and muscle artifacts. Ocular and additional artifacts were then corrected using independent component analysis (ICA). Next, data was segmented from 200ms pre-cue to 500ms post-cue onset (cue onset was determined using a photodiode), baseline-corrected (mean of the 200ms pre-cue subtracted from the data), and down-sampled to 200Hz. Our decision to limit the main analysis to the first 500ms post-cue onset was motivated by a need to ensure minimal influence of controlled retrieval processes following cue processing. Nonetheless, to ensure our results are not limited to this time windows, we repeated all MEG analysis for a longer time window (−200 to 800ms post cue onset; see Figure S4). All MEG analyses were performed at whole-brain sensor level.

#### Representation similarity semantic decoding

Semantic decoding was based on a recently developed representational similarity-based decoding method^37^. Semantic representations of cues were determined using three popular static embedding models, namely Word2vec^104^, GloVe^105^, and FastText^106^, and one contextualized embedding model, namely DistilBERT^53^. Whereas contextualized embedding models, which use a transformer architecture, are primarily used to quantify semantic representation of sentences or longer utterances (which is not necessary in a word association task), in our study we used this model to enable fine-tuning to specific individuals (see ‘Personalized semantic decoding’ below). We also tested semantic decodability using similarity metrics based on different transformations of free association norm data^51,52^, but these failed to show above-chance decodability.

Neural representation vectors were defined based on a cue-locked MEG response, separately for each time point. Representation similarity matrices were then calculated using Pearson correlation, whereas semantic similarity, in the baseline analysis, was averaged across the three static embedding models to reduce sensitivity to model-specific effects. Whereas standard representation similarity analysis correlates these matrices to calculate a representation similarity index, here we used a leave-2-out procedure to determine cue-level semantic decodability. Specifically, for every possible pair of cues used as the test-stimuli, the neural similarity vectors and the semantic similarity vectors are extracted. Importantly, the cues’ similarities to each other were removed to avoid overfitting. The decoding proceeds by a matching procedure: the decoder assigns ‘semantic’ labels by choosing the semantic similarity vectors that produce the best match to the neural similarity vectors (Figure 4A). Thus, successful classification is achieved if the sum of the diagonal elements of the neural-semantic correlation matrix is larger than the sum of the off-diagonal elements (Figure 1A).

The statistical significance of average decodability is determined by testing the null hypothesis of no relationship between semantic and neural similarity vectors. This null hypothesis is generated in a permutation test where the labels (i.e., rows and corresponding columns) of the semantic similarity matrix are shuffled, and are then correlated with the empirical neural similarity matrices^37^. Statistical significance was tested at the group level, and therefore the resulting decodability distributions under the null hypothesis were averaged across participants. Then, the significance threshold was determined, by first taking the maximum group-level null-decodability (across time points) to control for multiple comparisons, and then taking the 95% percentile of this distribution. These thresholds are depicted as horizontal lines in Figure 4. Next, to test the statistical significance of group differences, and the effects of speech disorganization symptoms, we used a cluster permutation test ^107^ based on the output of mixed-effects models, using the ‘permutes’ R package^108^. Here, we used a more conservative statistical threshold (0.01), while ignoring significant differences occurring earlier than 200ms, since we found no evidence for above-chance semantic decodability at these timepoints.

#### Personalized semantic decoding analysis

DistilBERT was fine-tuned using a bi-encoder architecture^53^, which can modify encoder weights to produce embeddings optimized for a given task. Specifically, we fine-tuned the model to align model-derived cosine similarity between each cue and the respective association with participants’ subjective similarity ratings, essentially encouraging the model to represent concepts in a way that reflects individuals’ semantic structures. To reduce over-fitting, we used a leave-one-out procedure, where the personalized embeddings of a given cue (and association) are predicted by a model trained on all other cue-association pairs. This also ensures that modified embedding of a given cue do not incorporate information regarding the associative retrieval processes it evoked. The resulting cue embeddings were then used to calculate personalized semantic RSM subsumed to our representational similarity decoding analysis depicted in Figure 4F. We used a cosine similarity loss, an AdamW optimizer with 100 warm-up steps, and either one, three or five epochs. As depicted in Figure S5, whereas using three epochs resulted in a considerable improvement in model performance, using five epochs did not improve performance, and, reduced decoding, suggesting over-fitting. Finally, we compared distilBERT to another, state-of-the-art, model: all-mpnet-base-V2. However, fine-tuning did not improve the performance of this, alternative model (Figure S6), rendering it unsuitable for obtaining personalized semantic representations.

