## Supplementary material for "The Semantic Underpinnings of Speech Disorganization in Schizophrenia": Figure S1

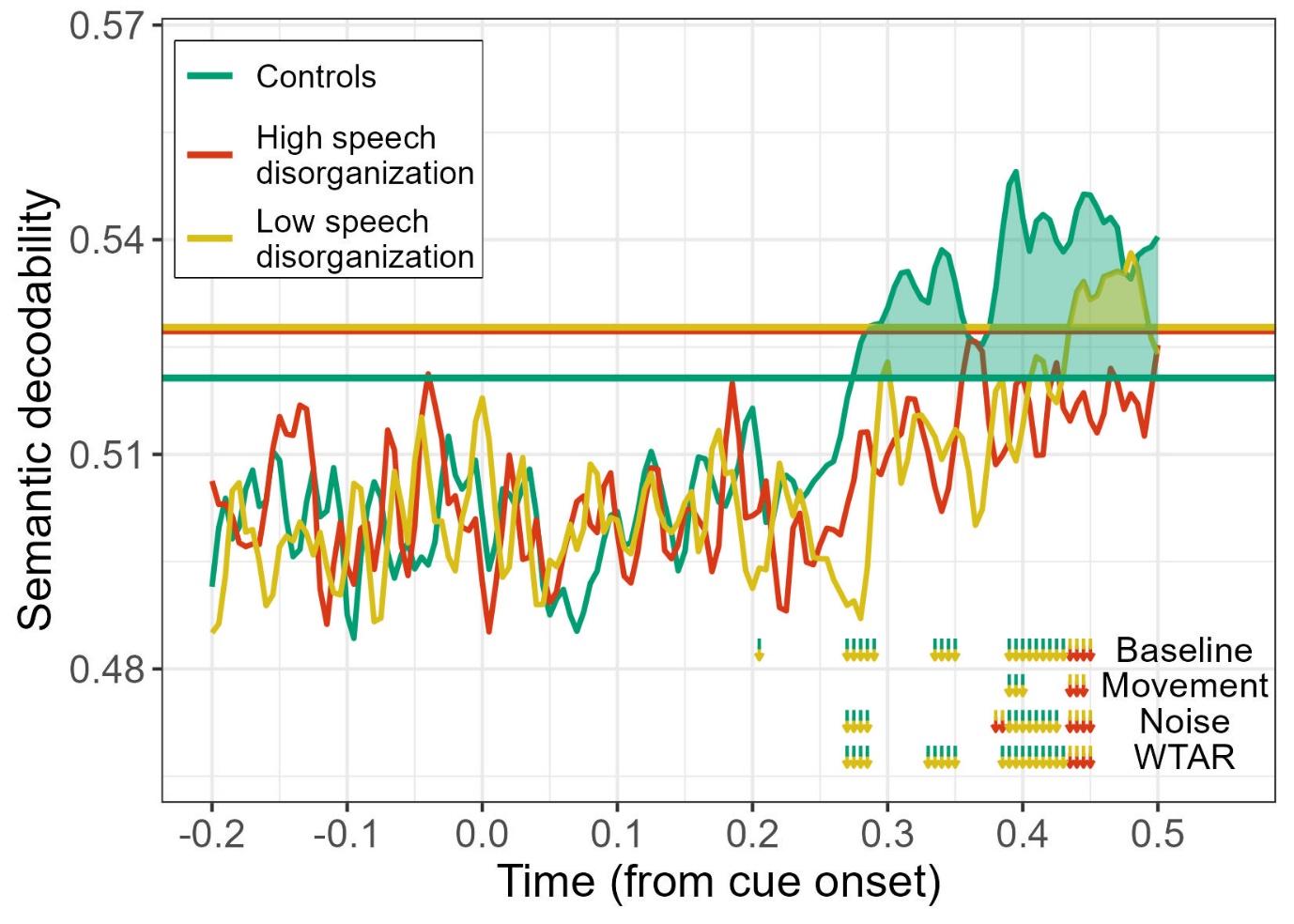

Figure S1 – Semantic decodability across groups controlling for several potential confounds (versus baseline model), particularly, head movements (Movement), neural noise (Noise), and verbal IQ (WTAR). Head movements were quantified combining two measures of head movements, namely, velocity, and the degree of displacement from the initial location. Both measures were averaged at the participant level. Neural noise was approximated using two method s. First, we examined variability inneural activity following the fixation point. Second, we examined variability of the neural representational similaritymatrix during the baseline period preceding cue onset.
