## Supplementary material for "The Semantic Underpinnings of Speech Disorganization in Schizophrenia": Figure S2

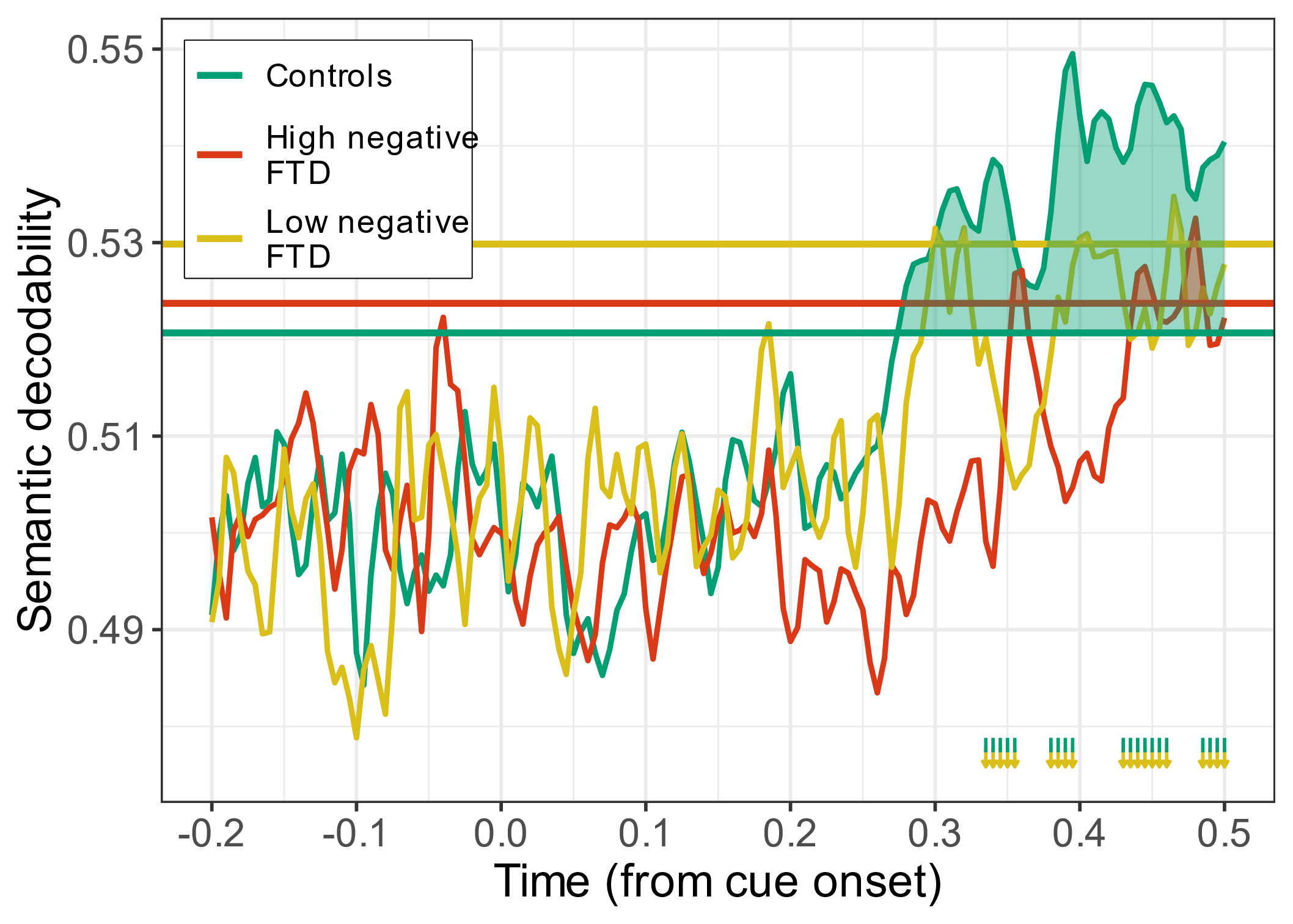

Figure S2 – Differences in semantic decodability when focusing on negative formal thought disorder (FTD) symptoms. We found no clear effects for negative FTD, supporting the specific role of speech disorganization (positive FTD) reported in the paper.
