## Supplementary material for "The Semantic Underpinnings of Speech Disorganization in Schizophrenia": Figure S3

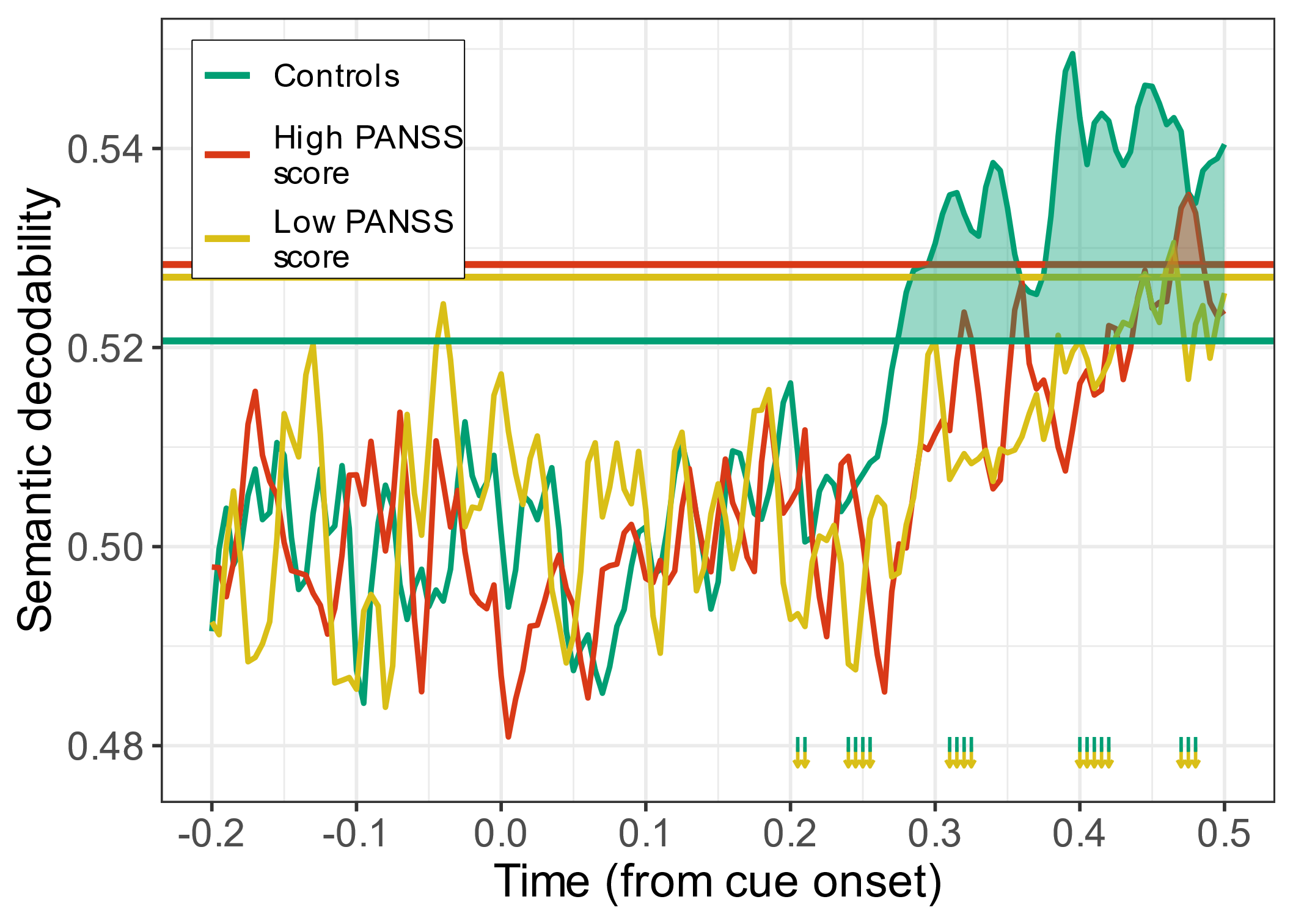

Figure S3 - Differences in semantic decodability as a function of general illness severity as measured by PANSS total scores (after excluding items pertaining to conceptual disorganization). We found no clear effects for general illness severity (and, in fact, some evidence for evident decodability among patients with high illness severity), providing further support for the specific role of speech disorganization (positive FTD) reported in the paper.
