## Supplementary material for "The Semantic Underpinnings of Speech Disorganization in Schizophrenia": Figure S4

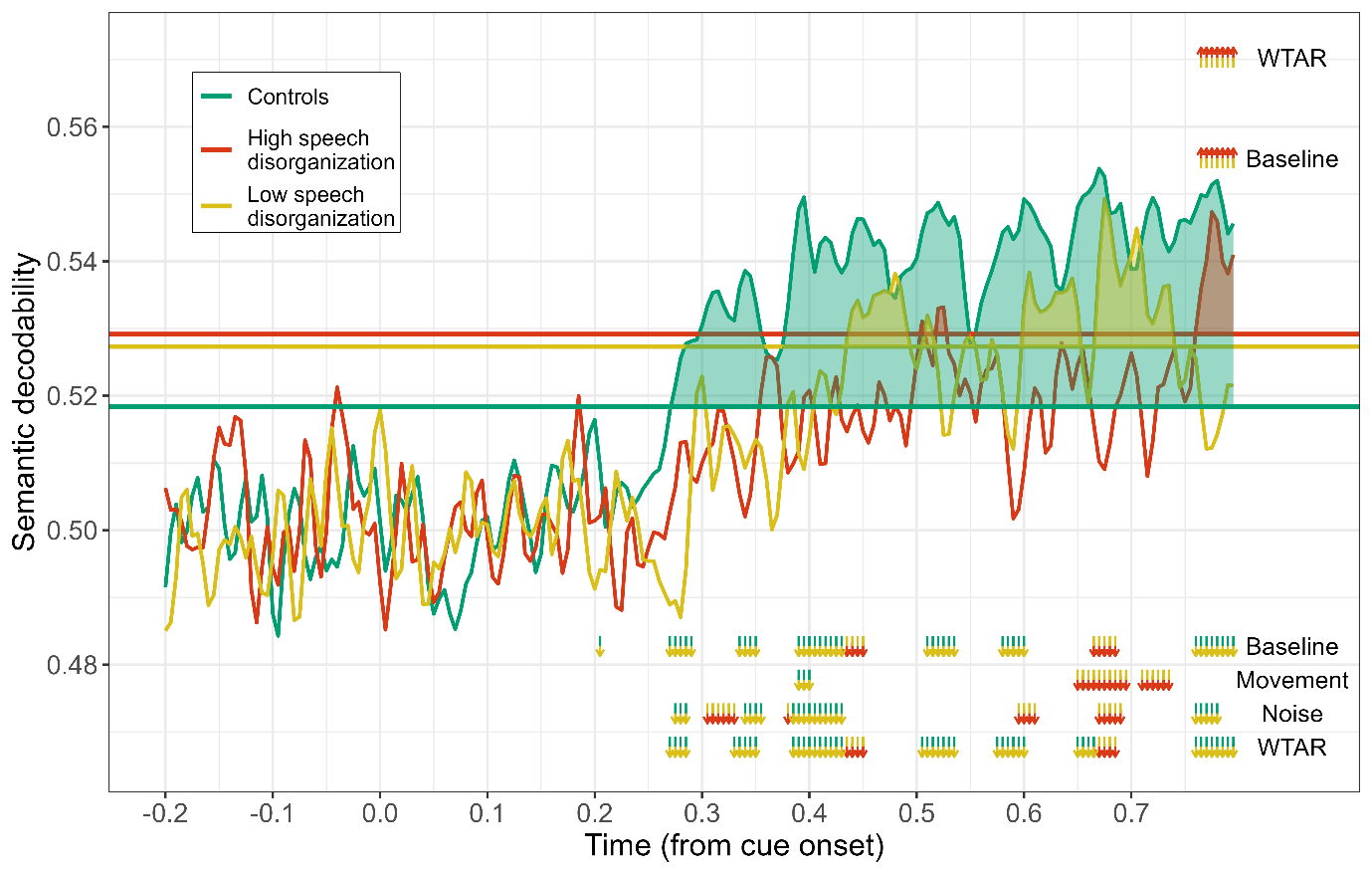

Figure S4 – Semantic decodability across groups measured for a longer time frame (-200-800ms). The results are generally consistent with the main findings reported in the paper but note an increase in decodability in those with speech disorganization starting around 700ms post cue onset. Crucially, however, this effect was no longer significant when controlling for either head movements or neural noise, further supporting the idea that the reduction in decodability in patients with low speech disorganization is partially explained by these confounds.
