## Supplementary material for "The Semantic Underpinnings of Speech Disorganization in Schizophrenia": Figure S5

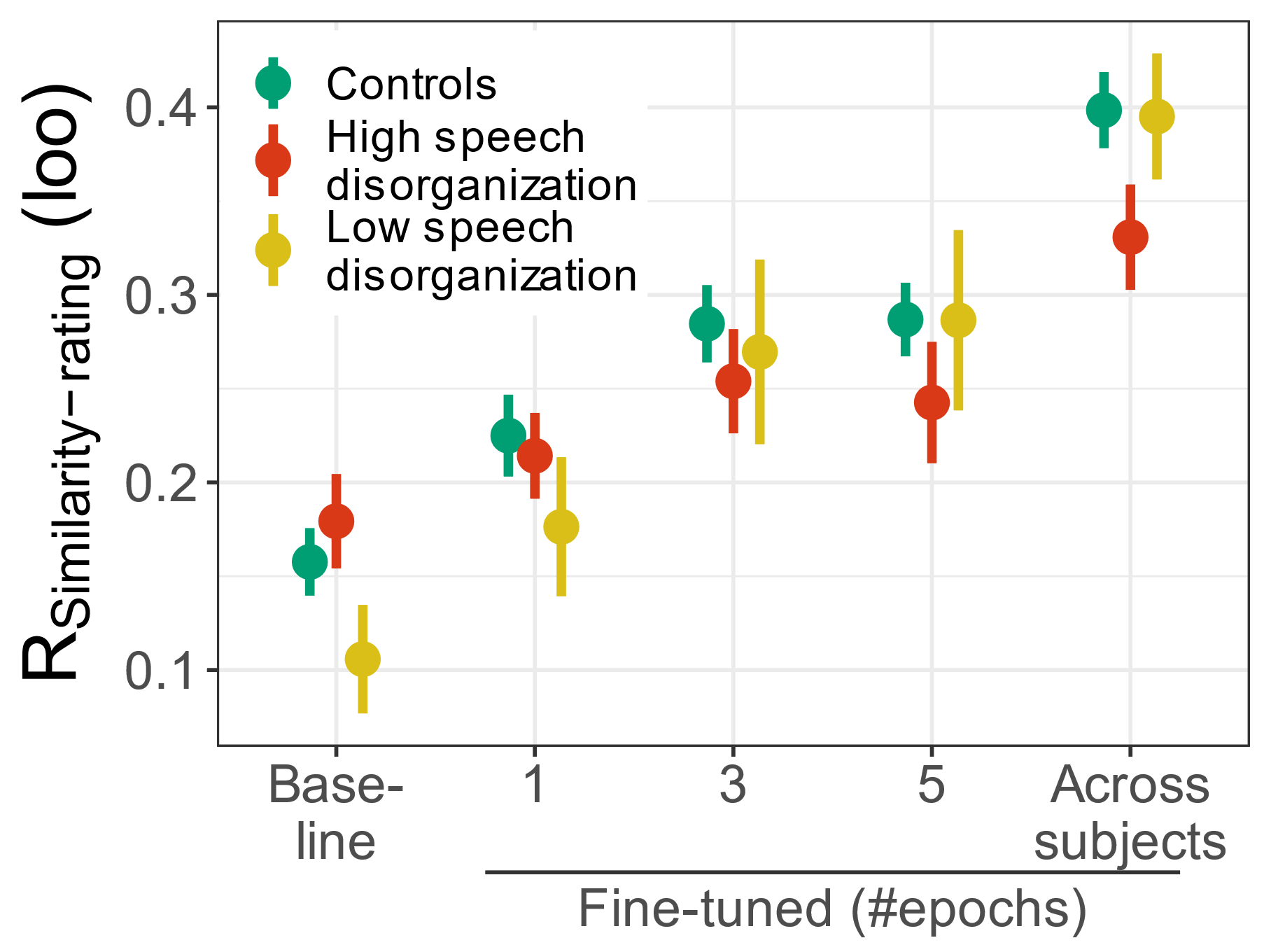
Figure S5 – Extended fine-tuning results. We tested fine-tuning using either one, three or five epochs. Whereas using three epochs resulted in a considerable improvement in model performance, using five epochs did not improve performance further. Thereby three epochs was used for the analyses reported in the main paper. Note also that groups did not differ in the effects of fine-tuning, except for patients with low speech disorganization symptoms, exhibiting a larger improvement following fine-tuning.
