## Supplementary material for "The Semantic Underpinnings of Speech Disorganization in Schizophrenia": Figure S6

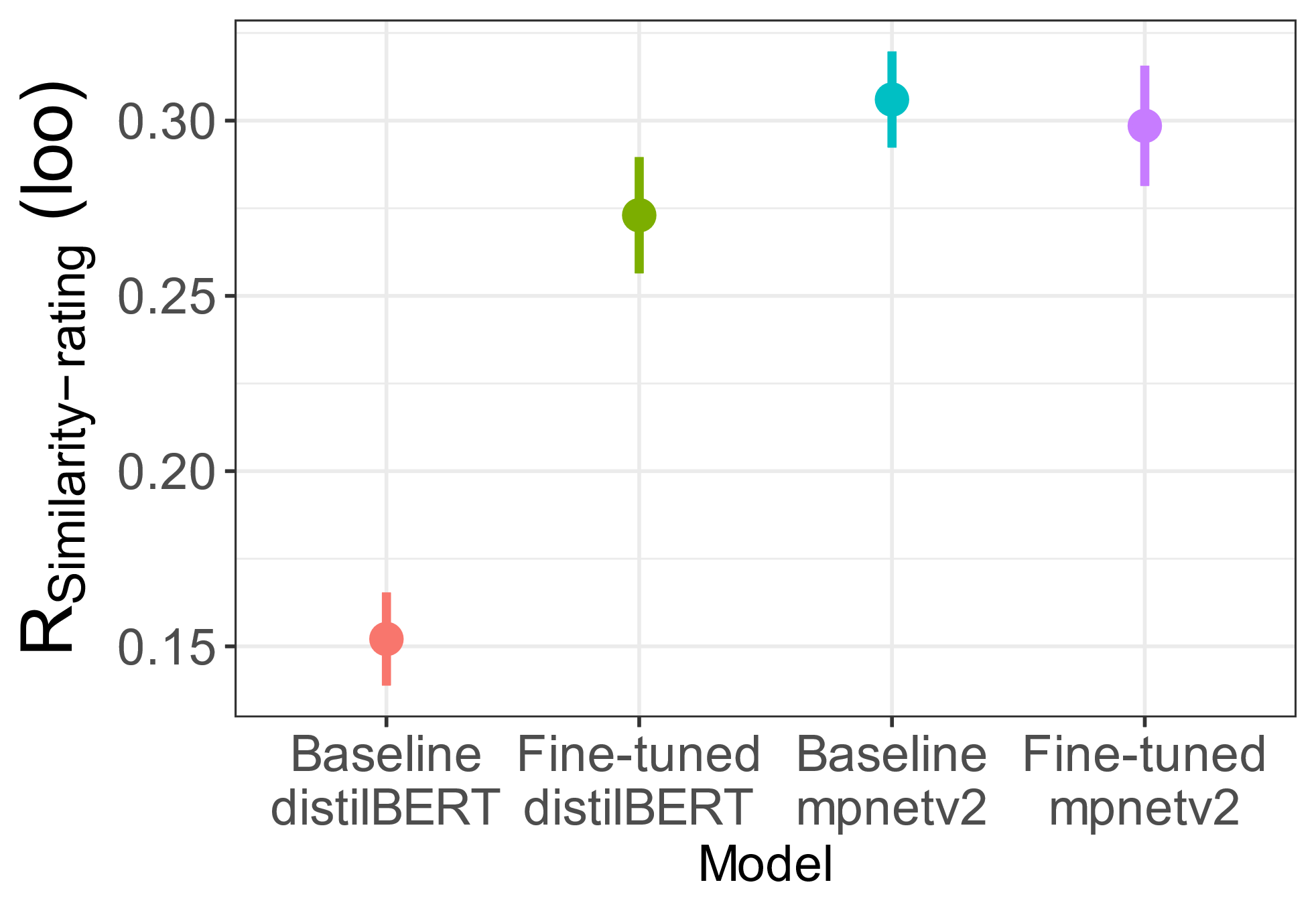

Figure S6 – Fine-tuning a different model. We tested whether using a different, state-of-the-art model (from the sentence-transformers library), namely *all-mpnet-base-V2* improves performance. Whereas this model had a higher correlation with associative strength ratings at baseline, fine-tuning did not further increase this correlation, suggesting that this model is not suitable for obtaining personalized semantic representations.
