## Supplementary Note S1 for "The Semantic Underpinnings of Speech Disorganization in Schizophrenia"

**Note S1 – Analyses using cosine similarity to measure atypicality of associations**

Typicality of associations can be measured either using word association norms, or continuous similarity measures extracted from language models, particularly word embedding models such as GloVE^[1](https://sciwheel.com/work/citation?ids=3581206&pre=&suf=&sa=0&dbf=0)^, Word2vec[^2^](https://sciwheel.com/work/citation?ids=13008707&pre=&suf=&sa=0&dbf=0), and FastText^[3](https://sciwheel.com/work/citation?ids=6853986&pre=&suf=&sa=0&dbf=0)^ We focus on the former in the main paper because we found norms based typicality measured to be more sensitive to our key manipulation. For comparison, here we report the key typicality-based results using cosine similarity averaged across the three key embedding models mentioned above. This analysis replicated the effect of speech disorganization symptoms on reducing typicality in the Free condition (*CI_Sdis|free_* = [-0.07, -0.006]; CI *_Sdis|restricted_* = [-0.05, 0.01]). Conversely, the effect of diagnosis on impaired the increase of typicality in the Restricted condition was not replicated (*CI_groupXcondition|Sdis=0_* = [-0.005, 0.003], *OR_condition|patients,Sdis=0_* = [0.001, 0.01], *OR_condition|controls,Sdis=0_* = [0.002,0.01]), potentially mirroring the generally weaker effect of the condition on cosine similarity.
