## Supplementary material for "The Semantic Underpinnings of Speech Disorganization in Schizophrenia": Table S1

| **Table S1 – Demographic and Clinical Variables** | | | |
| --- | --- | --- | --- |
| Variables | Controls (n=31) | Patients (n=31) | Statistical comparison |
| Gender |  |  | *χ^2^* = 1.03, *p* = .31 |
| Females | 13 (41.93%) | 18 (58.06%) |  |
| Males | 18 (58.06%) | 13 (41.93%) |  |
| Age | 32.10 (13.50) | 34.64 (9.62) | *t*(55.23) = 0.86, *p* =.4 |
| Education |  |  | *χ^2^* = 5.07, *p* = .28 |
| Lower secondary school | 1 (3.22%) | 0 (0.00%) |  |
| Upper secondary school | 11 (35.48%) | 10 (32.25%) |  |
| Level 4+ qualification | 4 (12.90%) | 10 (32.35%) |  |
| Bachelor’s degree or equivalent | 12 (38.71%) | 7 (22.58%) |  |
| Master’s degree or equivalent | 3 (9.68%) | 4 (6.45%) |  |
| Ethnicity |  |  | *χ^2^* = 5.85, *p* = .21 |
| White | 15 (48.38%) | 12 (38.71%) |  |
| African/Caribbean | 4 (12.90%) | 9 (29.03%) |  |
| East Asian | 1 (3.2%) | 1 (3.23%) |  |
| South Asian | 7 (22.58%) | 2 (6.45%) |  |
| Other/mixed | 4 (12.90%) | 7 (22.58%) |  |
| Country of primary education |  |  | *χ^2^* = 1.77, *p* = .18 |
| English speaking | 23 (74.19%) | 28 (90.32%) |  |
| Non-English speaking | 8 (25.81%) | 3 (9.67%) |  |
| WTAR (n = 59) | 107.75 (11.73) | 98.61 (14.13) | *t*(56.62) = -2.71, *p* =.008 |
| TALD (n = 60) |  |  |  |
| Speech disorganization (OP) | 0.08 (0.17) | 0.47 (0.47) | *t*(38.34) = 4.37, *p* <.001 |
| Reduced speech (ON) | 0.10 (0.20) | 0.87 (0.89) | *t*(33.08) = 4.71, *p* <.001 |
| Subjective positive | 0.58 (0.67) | 1.32 (1.14) | *t*(46.91) = 3.03, *p* =.003 |
| Subjective negative | 0.15 (0.16) | 1.00 (0.66) | *t*(33.76) = 6.88, *p <*.001 |
| MADRS (n = 60) | 15.94 (8.80) | 3.41 (3.58) | *t*(40.21) = 7.20s, *p <*.001 |
| PANSS (n = 30) | N/A | 61.77 (14.40) | N/A |
| Positive | N/A | 15.57 (5.59) | N/A |
| Negative | N/A | 14.40 (6.04) | N/A |
| General | N/A | 30.80 (6.37) | N/A |
| Primary diagnosis |  |  |  |
| Schizophrenia | N/A | 23 (74.19%) | N/A |
| Schizoaffective | N/A | 8 (25.81%) | N/A |
| Comorbidities | 0 (0.00%) | 10 (32.25%) | N/A |
| Panic disorder | 0 (0.00%) | 4 (12.90%) | N/A |
| Obsessive-compulsive disorder | 0 (0.00%) | 2 (6.45%) | N/A |
| Social anxiety disorder | 0 (0.00%) | 2 (6.45%) | N/A |
| Eating disorder | 0 (0.00%) | 1 (3.22%) | N/A |
| Substance use disorder | 0 (0.00%) | 3 (9.67%) | N/A |
| Age of Onset (24) | N/A | 26.71 (6.40) | N/A |
| Illness Duration (27) | N/A | 7.96 (7.52) | N/A |

**Note.** MADRS - Montgomery–Åsberg Depression Rating Scale; TALD - Thought and Language Disorder Scale; OP – Objective positive; ON – Objective negative; PANSS - Positive and Negative Syndrome Scale. Three patients reported an illness duration >25 years, but could not recall an exact age of onset.
